# Possible function of *Hox2* in atrial siphon fusion of the ascidian *Ciona*

**DOI:** 10.64898/2026.07.13.738359

**Authors:** Yurong Liu, Keita Yoshida, Akiko Hozumi, Kosuke Itageki, Nicholas Treen, Tetsushi Sakuma, Takashi Yamamoto, Toshinori Endo, Yasunori Sasakura

## Abstract

The hallmark of sessile adult ascidians is a vase-like shape with a single oral and atrial siphon. *Ciona*, however, develops two atrial siphons after metamorphosis, which subsequently fuse into one. The mechanisms underlying this fusion are unknown. This study suggests that *Hox2* controls this process. *Hox2*-knockout animals using Transcription-Activator-Like Effector Nuclease (TALEN) retain two atrial siphons throughout their lives. During normal fusion, epidermal cells between the siphons flatten along the anterior–posterior axis. This cellular flattening does not occur in *Hox2*-knockout animals, suggesting that the shape change in the epidermal cells produces tension, allowing the atrial siphon openings to converge at the midline for fusion. *Hox2*-knockout animals lack cupular organs, which are suspected hydrodynamic sensors in the internal epithelium of the fused atrial siphon and on the sperm duct. Among several knockout attempts, atrial siphon fusion was reproduced by only one TALEN pair, suggesting that this phenotype is driven by a mutation having a broader effect than those abolishing protein function. Many ascidians, unlike *Ciona*, develop a single atrial siphon shortly after metamorphosis. Our findings suggest that a phylogenetically conserved gene, *Hox2*, establishes this group-specific atrial siphon formation mechanism in *Ciona*.

## 1 INTRODUCTION

Ascidians are marine invertebrate chordates that constitute the largest subgroup within the subphylum Tunicata, and are closest living relative of Vertebrata (Satoh 1994; Delsuc et al. 2006). Ascidian larvae exhibit a typical tadpole shape and chordate characteristics such as a notochord and a dorsal neural tube. However, adult ascidians adopt a sessile lifestyle, unique among chordate groups. This adaptation results in body shapes markedly different from their larvae and vertebrates. All adult ascidians have a vase-shaped, with the pharynx and its gills comprising most of the body, adhering to a substrate at their base (Figure 1a). They feature an oral and an atrial siphon, protrusions that intake and excrete seawater, food, and digested materials (Figure 1a). While structures resembling ascidian siphons are absent in vertebrates, previous molecular and cellular studies suggest developmental association with vertebrate facial sensory systems (Kourakis & Smith 2007; Sasakura et al. 2012). The atrial siphon primordia (ASP) of ascidian larvae, visible as a thickened disk with a central opening on the trunk epidermis, express essential transcription factor genes for otic placode formation (Mazet et al. 2005; Wada et al. 1998). Adult atrial siphons possess cupular organs within their internal epithelium (Bone & Ryan 1978; Mackie & Burighel 2005; Mackie & Singla 2004; Millar 1953; Ohta et al. 2010). Cupular organs consist of sensory neurons and support cells, with a gelatinous matrix extending from the organ base into the atrial cavity. This morphology resembles fish lateral line organs and mammalian inner ear hair cells (Dijkgraaf 1964; Elepfandt 1988; Montgomery et al. 2000; Shelton 1971). These characteristics suggest the shared tunicate–vertebrate ancestor possessed a prototype auditory sensory system, which ascidians modified into the atrial siphon, possibly as an adaptation to sessility.

**FIGURE 1.**
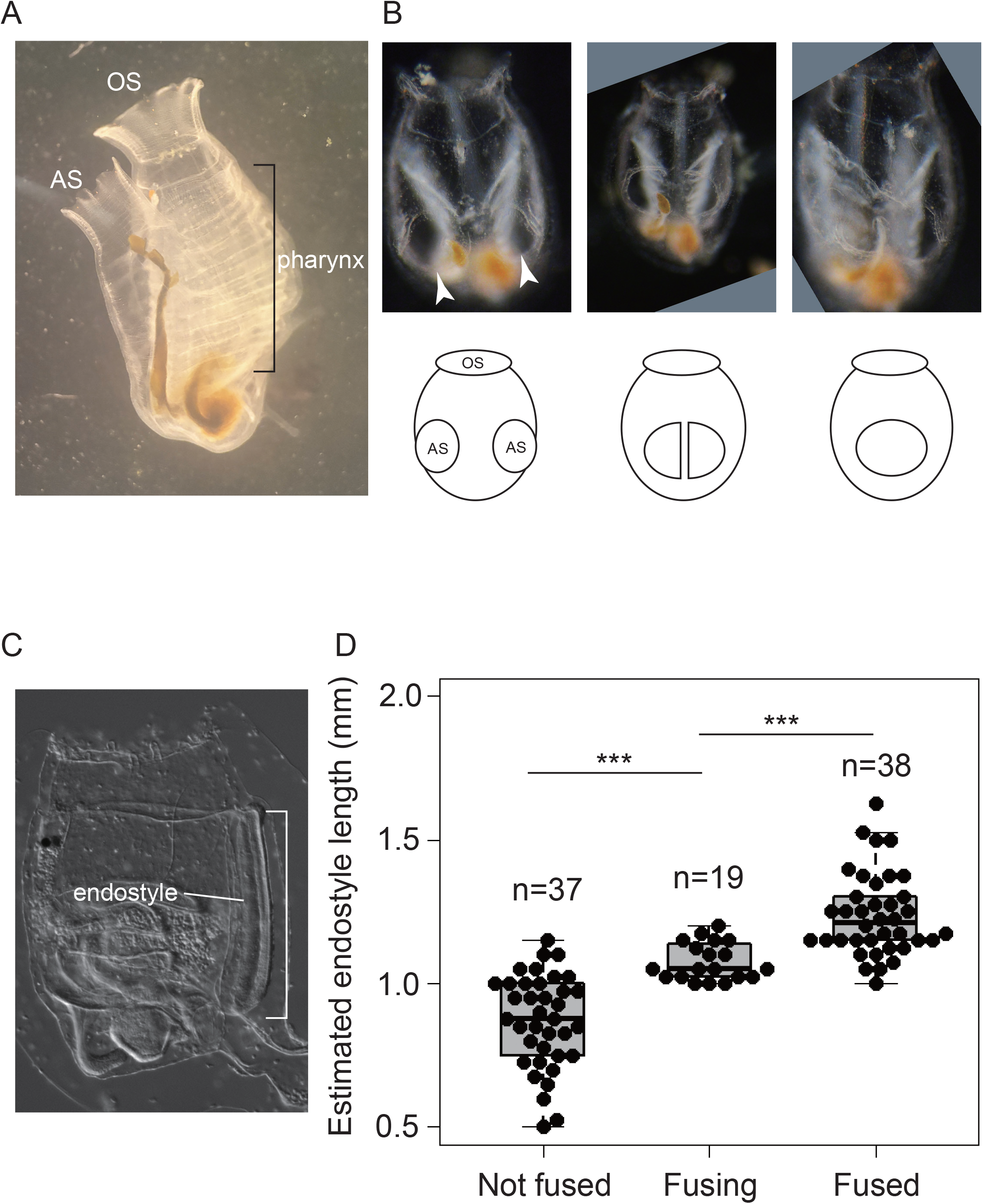
Atrial siphon fusion in *Ciona*. (a) *Ciona* adult after atrial siphon fusion. AS, atrial siphon; OS, oral siphon. (b) Atrial siphon fusion. The bottom illustrations are schematics based on the upper images. (c) *Ciona* juvenile showing the position of the endostyle. The bracket indicates how endostyle length was measured. (d) Relationship between endostyle length and atrial siphon fusion.

Adult ascidians, including the model species *Ciona intestinalis* Type A/*C. robusta* (Pennati et al. 2015), possess a single atrial siphon (Figure 1a). In contrast, *Ciona* larvae have two bilaterally paired ASPs (Kourakis & Smith 2007; Sasakura et al. 2012). This bilateral pairing may reflect a shared developmental and evolutionary origin with vertebrate ear placodes, which are also bilaterally paired structures. However, this hypothesis should be considered with caution, as many ascidian species have only one ASP (Satoh 1994). After metamorphosis, *Ciona*’s two larval ASPs develop into two atrial siphon openings in the juvenile stage, which subsequently fuse at the midline to form a single atrial siphon during growth (Figure 1b) (Berrill 1947; Ohta et al. 2010).

Our previous study, using enhancer-trap transgenic lines expressing GFP in cupular organ support cells, demonstrated that epithelia at the edges of the atrial siphon openings provide precursor cells for cupular organs (Ohta et al. 2010). During atrial siphon fusion, the midline edge epithelia migrate ventrally and subsequently form cupular organs. Thus, atrial siphon fusion is not merely a reduction in number; it is essential for establishing the atrial siphon’s internal sensory organs. Despite its importance, the molecular mechanisms underlying atrial siphon fusion remain poorly understood.

*Hox* genes encode homeodomain transcription factors critical for establishing characteristics along the anterior–posterior axis of metazoans (Gaunt 2018; Hueber & Lohmann 2008; Krumlauf 1994; Krumlauf 2018). These genes are typically arranged sequentially and tandemly in a single chromosom, commonly referred to as a cluster. A canonical *Hox* cluster in higher deuterostomes, including vertebrates, comprises 13 genes. *Hox* genes exhibit collinearity: a *Hox* gene located closer to the 3’ end of the chromosome, is expressed in a more anterior region and at an earlier developmental stage than one located closer to the 5’ end. Therefore, the chromosomal order of *Hox* genes correlates with their expression patterns and functions.

Ascidians possess a single, somewhat disorganized *Hox* cluster (DeBiasse et al. 2020; Dehal et al. 2002; Sekigami et al. 2017; Seo et al. 2004; Zhang et al. 2021). For example, *Ciona*’s cluster has a long gap between *Hox1* and *Hox2*; *Hox10* is located between *Hox4* and *Hox5*; *Hox7*‒*9* and *Hox11* are absent; and *Hox12* and *Hox13* reside on a different chromosome (Ikuta 2004). Despite this disorganization, *Ciona Hox* genes maintain spatial collinearity in their expression patterns within both the larval central nervous system and the adult digestive system (Ikuta et al. 2004; Nakayama et al. 2016). Similar disorganization in other tunicate species (Seo et al. 2004) suggests it predates the split of tunicate subgroups.

Ascidian larvae exhibit distinct anterior–posterior characteristics, including an anterior trunk and posterior tail. *Ciona Hox* genes are generally non-essential for establishing these larval anteroposterior characteristics (Ikuta et al. 2010). While *Hox* gene disruptions can lead to the loss of specific larval structures, such as ASP and a portion of neurons, larvae maintain a tadpole shape and complete metamorphosis. These observations suggest that *Hox* genes function primarily after embryogenesis. Indeed, genetic and genome-editing studies confirm that *Ciona Hox* genes play essential roles in adult body formation after metamorphosis (Kawai et al. 2015; Sasakura et al. 2012; Tajima et al. 2020; Yoshida, Nakahata et al. 2017). Specifically, *Hox1*, *Hox10*, and *Hox12* are crucial for digestive system development, and *Hox13* is required for the sensory organ at the sperm duct’s distal end. The adult ascidian body is increasingly providing insights into molecular mechanisms involved in forming the chordate body and for deducing the evolutionary history of chordates, because ascidian adults possess several chordate synapomorphies, such as the heart, gill, and endostyle (Diogo et al. 2015; Ogasawara et al.1999; Onuma et al. 2021; Yamagishi et al. 2022). Therefore, *Hox* genes are a primary research focus for understanding adult body formation in ascidians due to their central roles at this stage.

We used Transcription-Activator-Like Effector Nuclease (TALEN) to disrupt *Hox2* in *C. intestinalis* Type A (Cermak et al. 2011; Sakuma, Hosoi et al. 2013; Treen et al. 2014). We found that *Hox2* is likely essential for the fusion of the two atrial siphons during post-metamorphic growth; *Hox2*-disrupted animals developed into adults with two distinct atrial siphons, neither of which formed cupular organs. Many ascidian species possess only one atrial siphon and its primordia from larval to juvenile stages (Satoh 1994). Our study suggests that *Hox2*, a gene widely conserved among metazoans, mediates an event specific to a small group of ascidians, suggesting its role in speciation.

## 2 MATERIALS AND METHODS

### 2.1 Animals

Wild-type *C. intestinalis* Type A/*C. robusta* individuals, sourced from Onagawa Bay (Miyagi, Japan) and Onahama Bay (Fukushima, Japan), were cultivated as closed colonies by the National BioResource Project, Japan (Satou et al., 2026). Animals were maintained under constant light to prevent untimely spawning. Eggs and sperm were surgically collected from gonadal ducts. Epidermal GFP and enhancer trap lines for cupular organs were described previously (Ohta et al. 2010; Sasakura et al. 2010).

Animals were maintained using an in-land culture system (Joly et al. 2007) with recently modified conditions: seawater in tanks was exchanged weekly, and concentrated cell suspensions of *Pavlova sp.* and *Isochrysis sp.* (Reed Mariculture) were fed along with *Chaetoceros calcitrans*. Photographs were taken with an AxioImager Z1 and an AxioObserver Z1 (Carl Zeiss), or with an Interlens for iPhone 15 (Micronet) and an SZX16 objective microscope (Olympus).

### 2.2 Constructs

TALEN pairs targeting the *Hox2* homeodomain region were constructed on an *EF1α cis-*element and mCherry-tagged TALEN backbone vector using the Golden Gate method, as described previously (Cermak et al. 2011; Sakuma, Ochiai et al. 2013; Treen et al. 2014). Target sites were determined using TAL Effector Nucleotide Targeter 2.0 (Doyle et al. 2012). The *EF1α cis-*element was replaced with those of *TnI* (Davidson & Levine 2003), *AKR* (Hozumi et al. 2010), and *Epi-1* (Joly et al. 2007) using an In-Fusion Cloning Kit (Clontech), following prior reports (Sasakura et al. 2017; Yoshida, Hozumi et al. 2017; Yoshida & Treen 2018). TALEN repeat cDNAs were subcloned into pHTB>TALEN-NG::2A::mCherry for mRNA synthesis. TALEN mRNAs were synthesized *in vitro* using a MEGAscript T3 kit (Ambion), poly A tailing kit (Ambion), and cap structure analog (New England Biolabs). The *Hox2 cis*-element was isolated by PCR from *Ciona* genomic DNA using primers 5’-CGACTCTAGAGGATCCGTGTGTGAAAAACCAAGAG-3’ and 5’-GGCCGCAAGGGGATCCATCTCTGGATCCTGGCT-3’. This PCR fragment was inserted into the *Bam*HI site of pSPKaede (Horie et al. 2011) to create pSPHox2>Kaede. The PCR-amplified *Hox2*>*Kaede* cassette was then inserted into the *Bam*HI site of pMiLRneo (Klinakis et al. 2000) to create pMiHox2>Kaede. Table 1 lists the official construct names, following nomenclature rules (Stolfi et al. 2015). The guide RNA for *Hox2* knockout was custom-synthesized (Alt-R CRISPR-Cas9 crRNA) as previously reported (Sasakura & Horie 2023). The target sequence of the *Hox2* crRNA is 5’-TTACACCAACACCCAACTGCtgg-3’, with PAM shown in lowercase. Detailed methods are available at http://marinebio.nbrp.jp/ciona/.

**Table 1.** Official names of vectors and transgenic lines reported in this study.

| Abbreviated name or experimental aim in the manuscript | Full name according to the nomenclature rule |
| --- | --- |
| pSPHox2>Kaede | p*Ciinte*.REG.KY21.C1.14234077-14236112\|*Hox2*>*Kaede* |
| pMiHox2>Kaede | p(Mi-*Ciinte*.REG.KY21.C1.14234077-14236112\|*Hox2*>*Kaede*) |
| pBSEF1α>TALEN::2A::mCherry | p*Ciinte*.REG.KY21.C14.1351654-1353597\|*Eef1a.a*>TALEN::2A::mCherry |
| pBSEpi1>TALEN::2A::mCherry | p*Ciinte*.REG.KY21.C1.13658444-13660489\|*Epi-1*>TALEN::2A::mCherry |
| pBSAKR>TALEN::2A::mCherry | p*Ciinte*.REG.KY21.C1.12980740-12981761\|*AKR*>TALEN::2A::mCherry |
| pBSTnI>TALEN::2A::mCherry | p*Ciinte*.REG.KY21.C11.3012139-3013022\|*Tnni*>TALEN::2A::mCherry |
| *Hox2*>*Kaede* transgenic line or Tg[MiCiHox2K]1 | *Ciinte*.Tg[pMi-*Ciinte*.REG.KY21.C1.14234077-14236112\|*Hox2*>*Kaede*]1 |

### 2.3 Electroporation and microinjection

Unfertilized eggs were dechorionated with sodium thioglycolate and actinase E. Electroporation was performed as previously described (Corbo et al. 1997; Zeller 2018), using 60 μg per round. Microinjection into unfertilized eggs followed established protocols (Kobayashi & Satou 2018). Up to 500 μg/μl of TALEN mRNA was dissolved in microinjection solution containing 4 μg/μl FastGreen FCF (Fujifilm Wako). For CRISPR/Cas9-mediated knockout, 25 ng/μl of *Hox2* crRNA, 100 ng/μl of tracrRNA, and 400 ng/μl of Cas9 protein were dissolved in the microinjection solution, as previously reported (Sasakura & Horie 2023).

### 2.4 Transgenic and mutant lines

The Tg[MiCiHox2K]1 line, which expresses Kaede under the control of the *Hox2 cis*-element, was established via electroporation, as previously described (Matsuoka et al. 2005). *Hox2* knockout lines were generated using microinjection methods following prior reports (Sasakura & Horie 2023; Yoshida et al. 2014).

### 2.5 TALEN and CRISPR-Cas9 activity assays

TALEN activity was assessed as previously described (Treen et al. 2014; Yoshida & Treen 2018). *EF1α*>*TALEN::2A::mCherry* constructs were electroporated into one-cell embryos. Approximately 50 mCherry-expressing larvae were selected, and genomic DNA was isolated in bulk using a Wizard Genome DNA Isolation Kit (Promega). For *Hox2* guide RNA, genomic DNA was isolated from approximately 50 larvae injected with *Hox2* crRNA. The target sites of the corresponding TALENs and guide RNA were PCR-amplified using ExTaq DNA polymerase, Hot-Start version (Takara), and the amplified products were resolved on a 15% polyacrylamide gel according to a previous report (Ota et al. 2013). Primer sequences were: 5’-ATCGATAAGCTTGATGTTTTACGAGTTACCACATACG-3’ and 5’-CTGCAGGAATTCGATCCAAGAACCACATAATGCAGTG-3’ for TALENys and guide RNA, and 5’-CGCTTGTTACGTCAGAATCGTC-3’ and 5’-GTTTCCCATCGTGCCATGTAAG-3’ for TALENgl. PCR fragments exhibiting heteroduplex bands were subcloned, and mutations were determined by sequencing. Detailed methods are available at http://marinebio.nbrp.jp/ciona/.

To examine *Hox2* TALEN specificity, genomic DNA was isolated from four TALEN mRNA-injected and four uninjected animals using a Wizard Genome DNA Isolation Kit (Promega). Libraries were prepared using the TruSeq DNA PCR-Free Library Prep kit (Illumina) or, if that failed, the Nextera XT DNA Library Prep kit (Illumina). Genomic DNA was sequenced on a NovaSeq 6000 (Illumina) by Macrogen Japan.

### 2.6 Quality control, read mapping, and variant calling

Reads were trimmed for adapter sequences and quality-checked using fastp version 1.0.1 (Chen 2025) and fastQC version 0.12.1 (http://www.bioinformatics.babraham.ac.uk/projects/fastqc/). Quality-controlled reads were mapped to the *C. intestinalis* type A/*C. robusta* reference genome assembly (HT model; Satou et al. 2022) using BWA MEM version 0.7.17 (Li & Durbin 2009).

Mapped sequences were merged into a single BAM file using Samtools 1.13. Variant sites were identified using GATK tools version 4.5.0.0. The workflow involved: annotating duplicated reads with MarkDuplicates; identifying variations with HaplotypeCaller; integrating variation information across samples using GenomicsDBImport and GenotypeGVCFs; and applying hard filtering with SelectVariants and VariantFiltration. Finally, quality scores were recalibrated for sequence reads using BaseRecalibrator to obtain a BAM file, and the preceding steps were repeated.

### 2.7 Identification of candidate variations for siphon duplication

Variant calls were annotated for fidelity using SnpEff 5.3a, with an annotation database created from the HTmodel reference genome and the KY21 gene model. These annotations were evaluated using InterPro and a ColabFold-generated structural model to assess the functional impact of mutations. To identify TALEN off-targets, heterozygous variants were detected using Mutect2 and FilterMutectCalls (Benjamin et al. 2019; McKenna et al. 2010).

### 2.8 Genotyping

Genotyping followed the previously established method (Sasakura et al. 2003). To promote spawning, heterozygous mutant animals at the reproductive stage were dark-adapted for 30 min, then exposed to light. *Ciona* is hermaphroditic; progeny are obtained by self-fertilization, mixing eggs and sperm from a single individual for several hours. Fertilized eggs were collected in a 9-cm plastic dish and incubated at 18℃ until the larval stage. Post-metamorphosis, individual juveniles underwent digestion in 50 μl of TE containing 0.2 μg/μl proteinase K for 3 h at 50℃, followed by 15 min at 95℃ to inactivate the enzyme. One microliter of the digested solution was used as the PCR template. PCR conditions and primers matched those used in the TALEN activity assay. PCR fragments were subjected to 15% polyacrylamide gel electrophoresis and cloned for sequencing.

### 2.9 Phalloidin staining

Juveniles at the siphon fusion stage were fixed with 3.7% formaldehyde in seawater. Samples were rinsed several times in phosphate-buffered saline with 0.1% Tween20, 500 mM NaCl, and 100 mM Tris-Cl (pH 8.0) (PBSTNT). Specimens were then incubated with 2 units of Alexa Fluor 555-conjugated phalloidin (Invitrogen) dissolved in 200 µl PBSTNT for 1 h at room temperature. After multiple washes with PBST, specimens were observed using a confocal microscope (LSM700, Carl Zeiss).

## 3 RESULTS

### 3.1 Fusion of atrial siphons is size-dependent

Before examining *Hox2* function, we investigated whether the fusion of two atrial siphons was size- or age-dependent, thereby allowing us to determine the timing of this event more precisely. We empirically observed that fusion was size-dependent, as animals that exhibited severe growth retardation frequently had two atrial siphons, even when their siblings with larger body sizes had completed fusion. To quantify the relationship between body size and siphon fusion, we measured the length of the endostyle at the time of fusion. The endostyle is a straight, shape-stable organ that occupies a large part of *Ciona*’s body (Figure 1c); therefore, its length can be measured relatively precisely and used to compare body size among individuals (A. Nakayama et al. 2005). Animals that did not initiate siphon fusion had statistically shorter endostyle lengths than their siblings that had initiated or completed fusion (Figure 1d), confirming siphon fusion is size-dependent. The endostyle length of unfused animals did not exceed 1.5 mm. For our subsequent experiments, we observed atrial siphon fusion only after the animals had reached a sufficient size for this event to occur. Any animals that failed to reach this size were not analyzed.

### 3.2 A *Hox2* TALEN pair disrupts atrial siphon fusion

To investigate the role of *Hox2*, we generated a knockout using TALENs (Christian et al. 2010; Treen et al. 2014). We constructed a TALEN pair, named TALENys, targeting the homeodomain-encoding region. When expressed under a ubiquitous *EF1α* promoter, TALENys efficiently induced mutations at the target site (Figure 2a) (Sasakura et al. 2010, 2017; Treen et al. 2014). Since *EF1α*-driven TALEN expression can negatively affect embryogenesis due to its high levels, we introduced TALEN mRNAs to observe G0 generation phenotypes (Yoshida et al. 2014).

**FIGURE 2.**
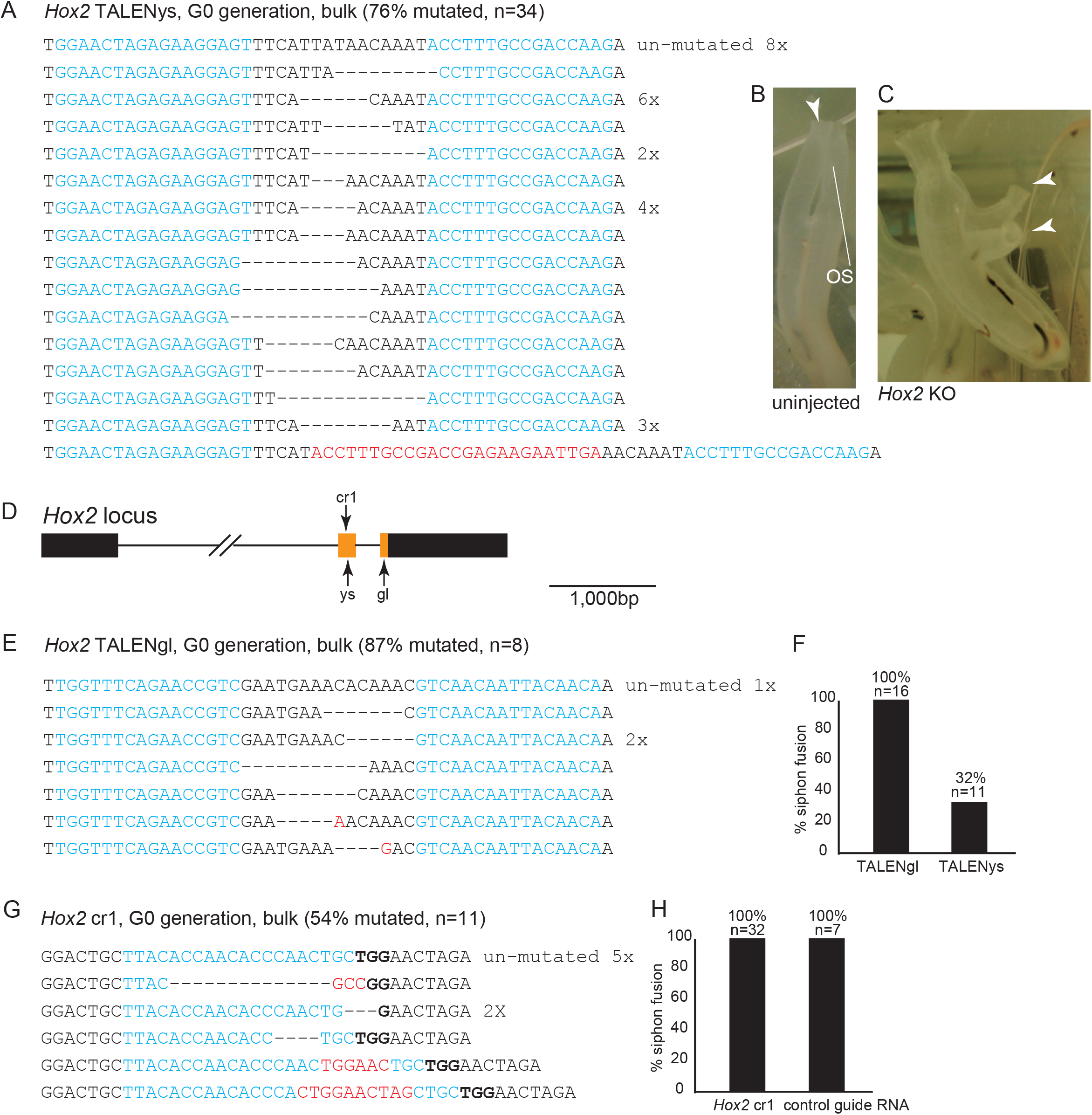
*Hox2* knockouts using TALENs and CRISPR/Cas9. (a) Examples of mutations induced by the pair of *Hox2* TALENys constructs in the G0 (TALEN-introduced) generation. TALEN binding sites are indicated in blue. The numbers on the right side indicate the number of clones obtained by sequencing. Bars represent deleted nucleotides. Red characters indicate inserted nucleotides. “Unmutated” indicates the reference sequence at the target site. (b) An uninjected control adult with one oral and one atrial siphon. The atrial siphon is shown by the arrowhead. (c) A G0 adult injected with *Hox2* TALENys mRNA. This animal has two atrial siphons, indicated by arrowheads. (d) Schematic illustration of the *Hox2* locus. Boxes and bars represent exons and introns, respectively. The region encoding a homeodomain is shown in orange. The target sites of TALENys, TALENgl, and the guide RNA are shown by the corresponding symbols. (e) Examples of mutations induced by the pair of TALENgl constructs in the G0 generation. (f) Atrial siphon fusion in G0 animals introduced with TALENgl or TALENys mRNA. (g) Examples of mutations induced by the guide RNA targeting *Hox2* (*Hox2* cr1) in the G0 generation. The PAM sequence is shown in bold. (h) Atrial siphon fusion in G0 animals introduced with *Hox2* cr1 or a control guide RNA. The guide RNA targeting *tyrosinase* was used as the control.

The animals in which *Hox2* was knocked out using TALENys mRNA exhibited normal embryogenesis and developed into normal larvae. They underwent normal metamorphosis and became juveniles that were indistinguishable from wild-type animals. However, the *Hox2*-knockout animals did not fuse their two atrial siphons, even after reaching a size sufficient for fusion to occur (Figure 2b and c). As a result, *Hox2*-knockout animals retained two atrial siphons throughout their lives. We termed this abnormality the “two-atrial-siphon” phenotype.

To confirm *Hox2* as the causative gene for this phenotype, we attempted knockout with a different TALEN pair (TALENgl), targeting a distinct DNA sequence (Figure 2d,e). Surprisingly, TALENgl did not induce the two-atrial-siphon phenotype (Figure 2f), despite its higher mutation efficacy compared to TALENys. We also performed *Hox2* knockout using CRISPR-Cas9 (Jinek et al. 2012; Sasaki et al. 2014). One guide RNA (cr1; Figures 2d and 2g) showed moderate mutation efficiency, but animals with cr1-induced *Hox2* knockout did not exhibit the two-atrial-siphon phenotype in the G0 generation (Figure 2h). To further demonstrate that *Hox2* cr1 mutations did not impair atrial siphon fusion, we generated homozygous *Hox2* mutants. Crossing two G0 individuals with a high likelihood of germline *Hox2* mutations (G0-1 x G0-2 in Figure 3a) yielded no progeny with the two-siphon phenotype (0%, n=22).

**FIGURE 3.**
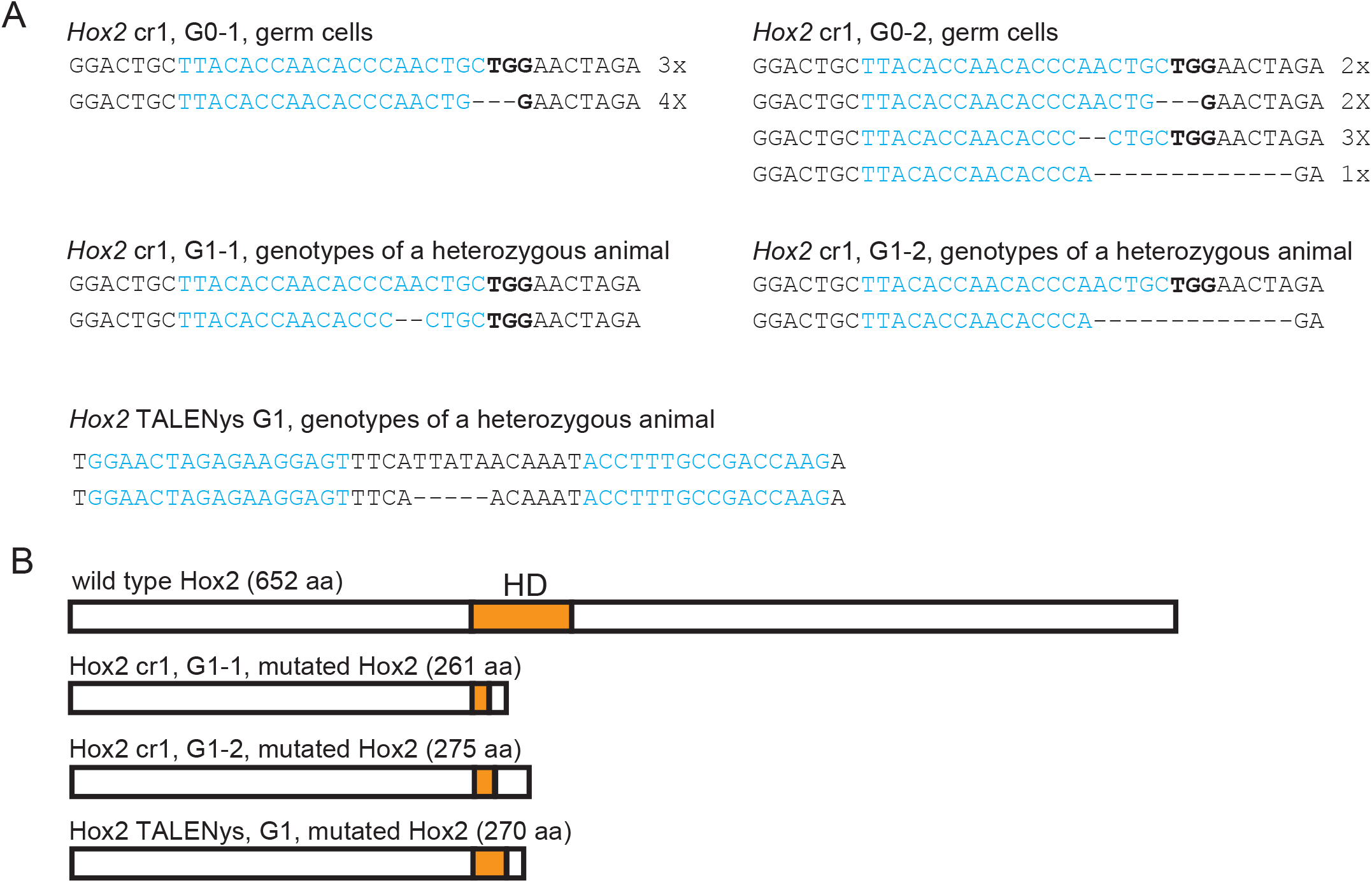
*Hox2* homozygous mutants did not exhibit the two-atrial-siphon phenotype. (a) Mutation patterns detected in the corresponding samples. (b) Deduced structures of the Hox2 protein translated from the mutated alleles shown in (a). HD, homeodomain; aa, amino acid.

Similarly, a cross of two *Hox2* heterozygous mutant animals (G1-1 x G1-2 in Figure 3a) produced no affected progeny (0%, n=13). The mutated *Hox2* alleles in G1-1 and G1-2 were predicted to cause frameshifts, resulting in truncated Hox2 proteins lacking most of the homeodomain (Figure 3b). To examine whether a TALENys-induced mutation could yield the two-atrial-siphon phenotype, we established a mutant line harboring a 5-nucleotide deletion in the homeodomain-coding region, induced by TALENys (*Hox2* TALENys G1, Figure 3a). Homozygous mutants from this line did not exhibit the abnormality (0%, n=3). Thus, only transient *Hox2* mutations induced by TALENys reproducibly caused the two-atrial-siphon phenotype.

These results suggest that the two-atrial-siphon phenotype cannot be explained by simple loss of *Hox2* function. One possible explanation is that TALENys induced mutations in another gene through an off-target mechanism. However, such off-target effects are less probable with TALENs than with CRISPR-Cas9, as TALEN pairs target a relatively long DNA stretch (32‒34 nucleotides in our standard protocol; Sakuma, Ochiai et al. 2013; Treen et al. 2014). Indeed, a blastn survey of the *Ciona* genome revealed no potential off-target sites with sufficient nucleotide sequence conservation to the TALENys target site. To confirm that the *Hox2* homeodomain-encoding site was the only region mutated by the TALENys pair, we sequenced genomic DNA from somatic cells of four TALENys-introduced animals and four uninjected controls (Table 2). After individually assembling the genome sequences, we identified mutations present exclusively in the TALENys-introduced animals. This analysis revealed 6,758 such sites, far exceeding expectations (Figure 4a). We then screened these sites for features consistent with TALEN-induced mutations in the G0 generation (Figure 4a):

1. **Mosaicism:** Several patterns of TALEN-induced mutations were detected in the genomes isolated from the whole body at the G0 generation because two blastomeres may be mutated independently if they divide before mutation introduction.
2. **Target-site location:** TALEN-induced mutations occur in the spacer between the binding sites of two TALENs. Because our TALENs recognize 16‒17 nucleotides, which are separated by a 15-nucleotide spacer, TALEN-induced mutations should accumulate within a 48-nucleotide region.
3. **Exclusion of misassemblies:** Misassemblies of the genome sequence must be considered and excluded from the analysis. These misassemblies tend to occur in repetitive sequences.
4. **Deletion bias:** TALENys-induced mutations tend to be multi-nucleotide deletions (Figure 2a). In other words, single-nucleotide variations are more likely to be due to sequencing errors.
5. **Absence from wild-type populations:** Because the two-atrial-siphon phenotype is specific to TALENys-introduced groups, the causative mutations induced by TALENys should not be present in the genomes of the wild-type populations used in this study. The resource project in Japan sequenced the genomes of wild-type populations (Satou et al., 2026), which can be retrieved from the Ghost genome database (http://ghost.zool.kyoto-u.ac.jp/default_ht.html).

**FIGURE 4.**
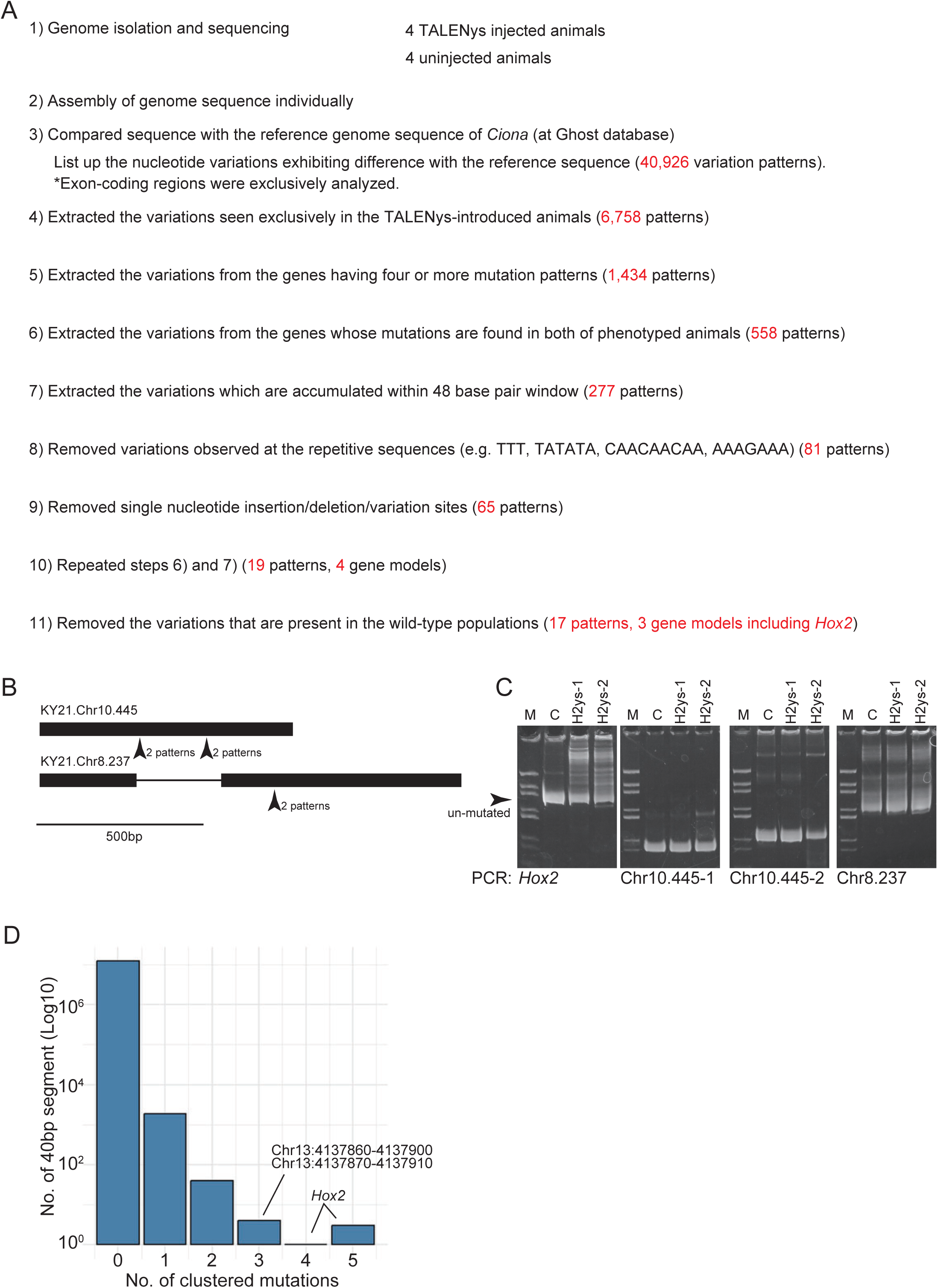
*Hox2* is the only gene mutated efficiently by TALENys. (a) Step-by-step procedure used to analyze the variation sites and patterns found in the sequenced genomes. (b) Variation sites remaining after the screening in (a). Their positions in the gene models are shown by arrowheads. Each position includes two variation patterns. Variations in *Hox2* are not shown in this panel. Boxes and bars represent exons and introns, respectively. (c) *Hox2* is the only gene with reproducibly detectable sequence variations in the genomes of TALENys-introduced animals. M, marker lane (pBluescript digested with *Hae*III); C, control lane (an animal introduced with a TALEN pair unrelated to *Hox2*); H2ys-1 and −2, two *Hox2* TALENys-introduced animals that were not used in the analysis in (a). In the left-side gel image showing the *Hox2* analysis, higher molecular weight bands appear in H2ys-1 and H2ys-2 lanes in addition to the major band, which corresponds to unmutated *Hox2* (arrowhead). These additional bands represent sequence variations, indicating mutations induced by *Hox2* TALENys. In the analyses of the other genes, there is no difference between control and *Hox2* TALENys-introduced animals, indicating that TALENys did not induce mutations in these genes. (d) Number of clustered mutation sites in 40-bp segments. The X and Y axes correspond, respectively, to the number of observed clustered mutations and the number of segments that contain them.

**Table 2.** Sequence information for genomes of *Hox2* TALENys-introduced and control animals.

| Sample | Library | Total sequences<br>(million bp) | Mapped reads<br>(%) | Mean depth<br>(x) | Genome coverage<br>($\geq 1x$, %) |
| --- | --- | --- | --- | --- | --- |
| TALENys-1 | Nextera | 14.6 | 98.4 | 16.6 | 97.0 |
| TALENys-2 | Nextera | 38.6 | 99.0 | 45.5 | 99.0 |
| TALENys-3 | TruSeq | 77.2 | 99.1 | 90.8 | 99.0 |
| TALENys-4 | TruSeq | 78.1 | 99.0 | 91.6 | 99.0 |
| Control-1 | TruSeq | 81.5 | 98.9 | 95.5 | 99.0 |
| Control-2 | TruSeq | 77.9 | 99.0 | 91.5 | 99.0 |
| Control-3 | TruSeq | 80.7 | 99.0 | 94.7 | 99.0 |
| Control-4 | Nextera | 37.5 | 98.9 | 44.2 | 98.0 |

We analyzed only the genomic regions having a gene model (Satou et al. 2022) and found 17 variations that met our criteria (Figure 4a and Table 3). Eleven were in the *Hox2* TALENys target site. The remaining six were in two gene models (KY21.Chr10.445 and KY21.Chr8.237) (Figure 4b). Neither gene has a substantial open reading frame, and neither contains coding-region sequences homologous to the TALENys-binding sites. We then tested for reproducibility of these mutations in *Hox2* TALENys-introduced animals not used for initial genome sequencing. No mutations were detected in KY21.Chr10.445 or KY21.Chr8.237 in two samples (Figure 4c), indicating that the mutations in these genes were not reproducible and were therefore unlikely to be causative of the two-atrial-siphon phenotype.

**Table 3.**
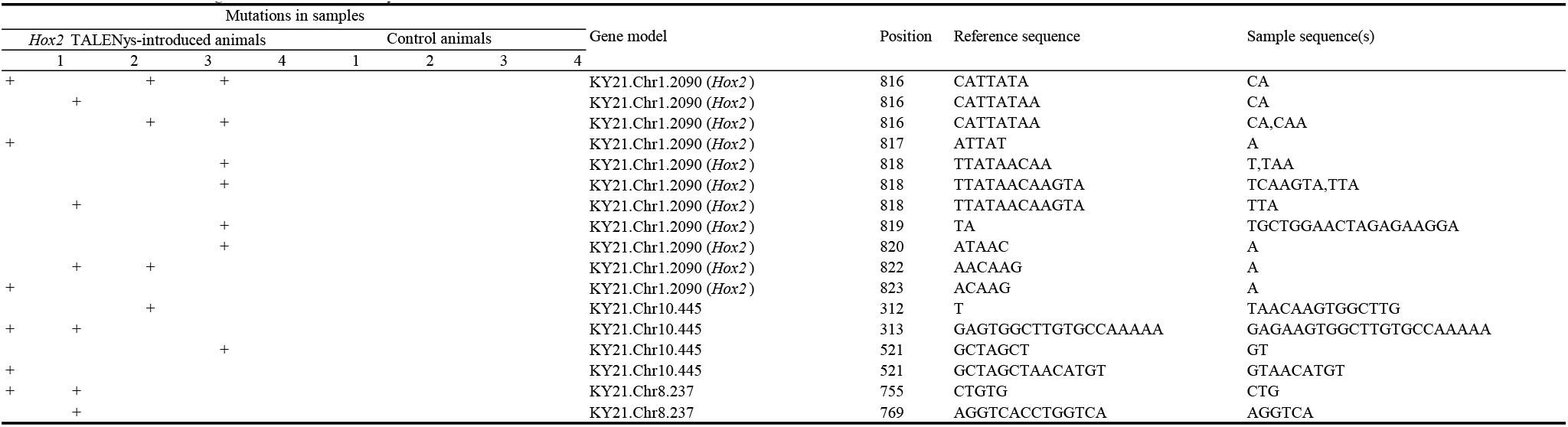
Mutations found in the genomes of Hox2 TALENys-introduced G0 animals.

While the initial analysis focused on coding regions, a causative mutation might reside in a non-coding region. To investigate this, we performed a genome-wide survey of TALEN-mutated sites using TruSeq data with sufficient read depth. Assembled sequences were segmented into 40-nucleotide stretches with 30-bp overlaps, and clustered mutations were counted per segment (Figure 4d). The segments encompassing the *Hox2* TALENys-binding site showed the highest number of clustered mutations, confirming the specificity of TALENys. The third-highest number of clustered mutations occurred in a region on chromosome 13 (two segments between 4,137,860 and 4,137,910). This region, an intron of gene model KY21.Chr13.462, is rich in GC (5’-AGAGGGGGTGGGGGTTAGAGGGGGGGTGGGTTGGATGGGCGGTTAGGGGGTT-3’), suggesting that the observed mutations are due to misassembly. We concluded that the *Hox2* locus was the only reproducibly mutated candidate associated with the two-atrial-siphon phenotype. Subsequent experiments used TALENys to perturb *Hox2* function for functional analyses.

### 3.3 Possible *Hox2* role in the epidermis to promote atrial siphon fusion

The mechanism for atrial siphon fusion are unknown. To gain insight into this process, we knocked out *Hox2* in a tissue-specific manner by overexpressing TALENys under the control of tissue-specific *cis*-elements. We selected the epidermis, muscle, and mesenchyme for this experiment. The two-atrial-siphon phenotype was reproduced only when TALENys was expressed in the epidermis (Figures 5a-c). This result suggests that the epidermis is responsible for the fusion of the siphons.

**FIGURE 5.**
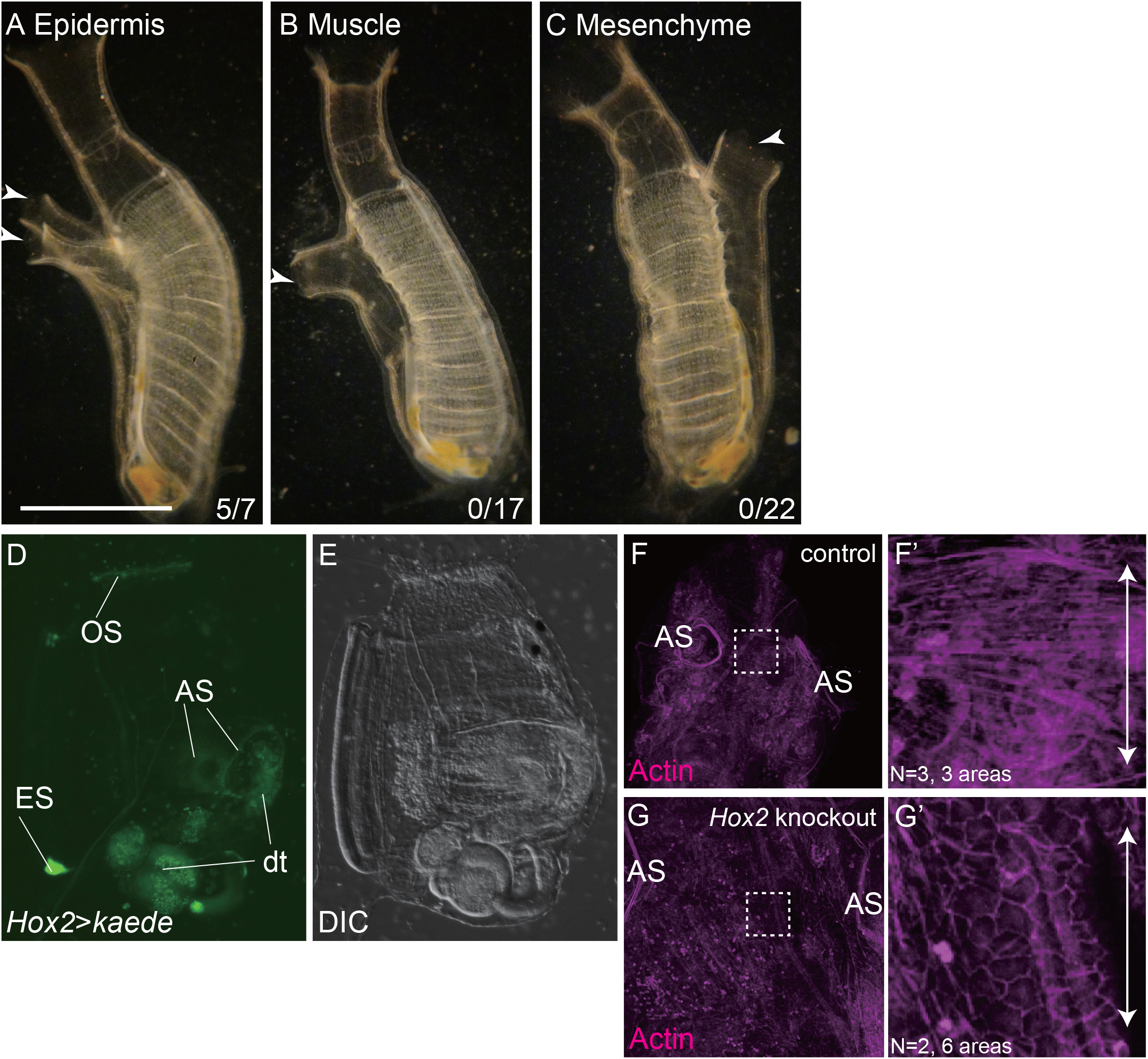
*Hox2* in the epidermis is likely responsible for atrial siphon fusion. (a) A G0 animal expressing *Hox2*-targeting TALENys in the epidermis, with two atrial siphons (arrowheads). The number at the bottom represents the number of animals exhibiting the two-atrial-siphon phenotype. Bar, 5 mm. (b) A G0 animal expressing *Hox2*-targeting TALENys in the muscle, with one atrial siphon (arrowhead). (c) A G0 animal expressing *Hox2*-targeting TALENys in the mesenchyme, with one atrial siphon (arrowhead). (d-e) Kaede expression in the *Hox2*>*Kaede* transgenic line at the juvenile stage before atrial siphon fusion. AS, atrial siphon; dt, autofluorescence in the digestive tube; ES, the posterior end of the endostyle; OS, oral siphon. (e) is the differential contrast image of (d). (f-g) Shape of epidermal cells between two atrial siphons during their fusion, observed by staining actin filaments with phalloidin. (f-f’) A control animal. (f’) is an enlarged image of the area shown by the dotted line in (f). The arrows indicate the orientation of the anterior–posterior axis. (g-g’) A *Hox2* TALENys-introduced animal.

S. Nakayama et al. (2016) described the expression pattern of *Hox2* in detail using whole-mount *in situ* hybridization at the early adult stage, soon after siphon fusion. Although those authors did not focus on epidermal expression, their published image suggested *Hox2* expression in the atrial siphon epidermis (Fig. 1c in Nakayama et al., 2016). To confirm *Hox2* expression in the epidermis, we established a transgenic line harboring a reporter construct containing a *Hox2 cis*-element. This transgenic line expressed Kaede fluorescent protein in the epidermis around the atrial siphon openings before their fusion (Figure 5d-e). This line also showed Kaede expression at the oral siphon edge and the posterior end of the endostyle. Because S. Nakayama et al. (2016) showed that *Hox2* is strongly expressed in the posterior end of the endostyle, this finding suggests that our reporter construct recapitulates the endogenous expression pattern of *Hox2*.

We observed the shape of epidermal cells between the two atrial siphons as they converged. In control animals, epidermal cells flattened along the anterior–posterior axis when viewed apically (Figures 5f and f’). Conversely, *Hox2* TALENys-introduced animals at the same age did not flatten, with epidermal cells retaining their polygonal shape (Figures 5g and g’). This suggests that changes in epidermal cell shape drive siphon fusion.

### 3.4 *Hox2* TALENys disrupts cupular organ formation in the atrial siphon

Our previous study suggested that cupular organs on the atrial siphon epithelium and sperm duct form through siphon fusion (Ohta et al. 2010). During the fusion of two atrial siphons, the joined siphon edges migrate ventrally along the midline (Figure 6a). Cupular organs emerge from these migrating edges, which eventually fuse to the branchial epithelium, forming the wall of the sperm duct. This observation explains why many cupular organs align along the duct (Figure 6b). We hypothesized that *Hox2* knockout with TALENys would result in the absence of cupular organs in the atrial epithelium and sperm duct due to failed atrial siphon fusion. To test this, we introduced TALENys into a cupular organ marker transgenic line (Ohta et al. 2010). As predicted, cupular organs were absent in both the atrial epithelium and sperm duct of *Hox2* TALENys introduced animals (Figure 6c). Our cupular organ marker line, created by enhancer trap (Awazu et al. 1987), expresses GFP in both cupular organs and the orange pigment organ (OPO) at the anterior edge of the sperm duct (Figure 6b; Ohta et al. 2010), suggesting shared developmental mechanisms. However, in *Hox2* TALENys introduced animals, the OPO still exhibited GFP expression (Figure 6c), suggesting that the role of *Hox2* is limited to cupular organ formation.

**FIGURE 6.**
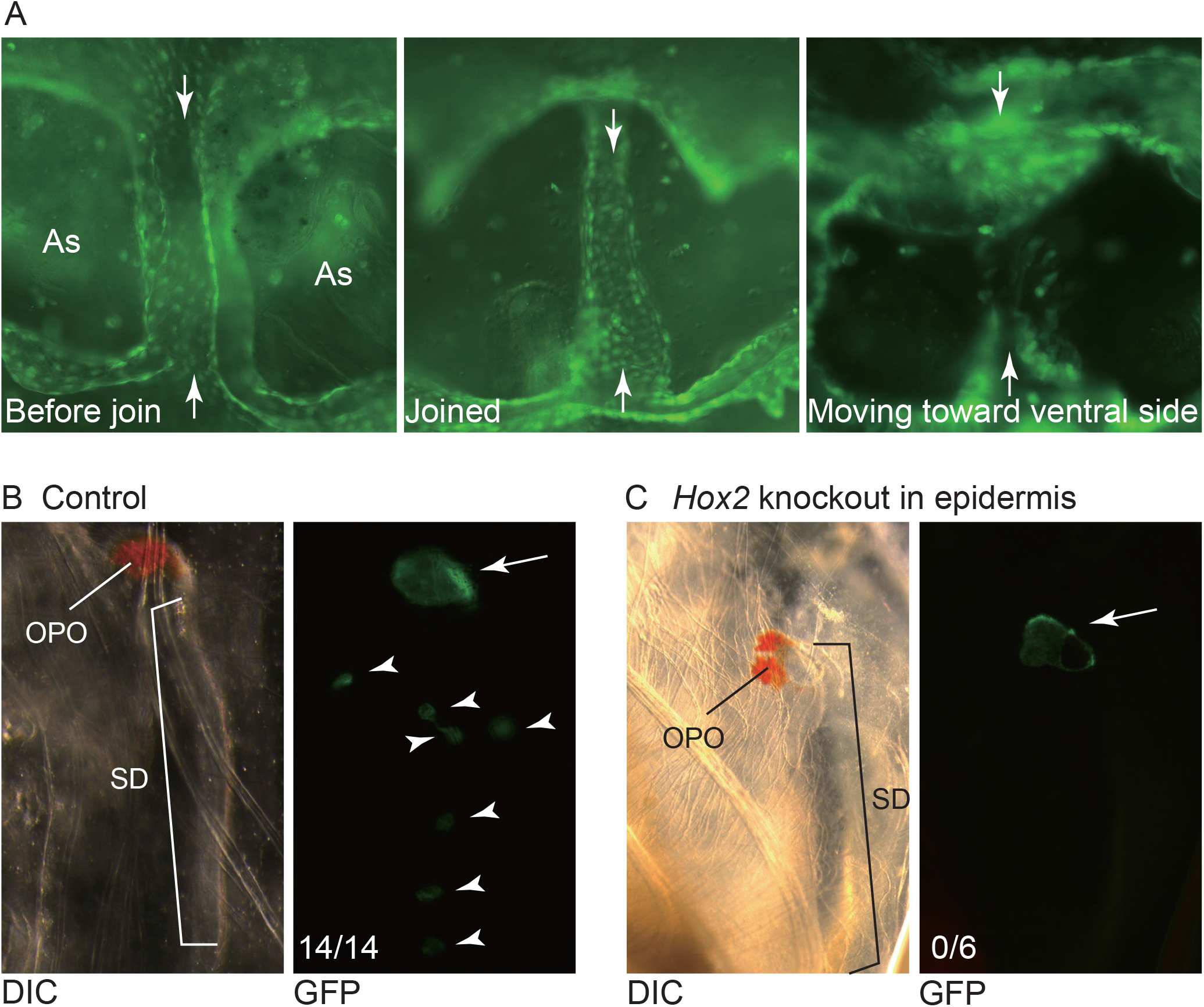
Atrial siphon fusion is required for cupular organ formation. (a) Fluorescent images of an epidermal GFP transgenic line showing migration of the midline epithelia of the two atrial siphons (As). The midline is indicated by arrowheads. (b) The sperm duct (SD) of a control adult with GFP expression in the cupular organs (arrowheads) and an orange pigment organ (OPO) at the terminus of the sperm duct (arrow). (c) *Hox2* TALENys-introduced adult that does not exhibit GFP expression corresponding to cupular organs. The OPO exhibited GFP fluorescence.

## 4 DISCUSSION

*Hox* genes are essential for establishing anterior–posterior axis characteristics in animals (Hueber & Lohmann 2008). The tadpole shape, with an anterior trunk and posterior tail, is a shared feature of higher chordates or olfactants. In vertebrates, *Hox* genes play crucial roles in embryogenesis (Krumlauf 1994; Mallo 2018). However, ascidian *Hox* genes are unnecessary for tadpole larval body formation (Ikuta et al. 2010), possibly due to their highly simplified larval body. In contrast, our previous studies show that Ciona’s Hox genes are essential for juvenile body formation after metamorphosis (Kawai et al. 2015; Sasakura et al. 2012; Tajima et al. 2020; Yoshida, Hozumi et al. 2017; Yoshida, Nakahata et al. 2017). This study suggests that *Hox2* meets this criterion. After metamorphosis, ascidians lose their tadpole shape, becoming vase-like adults. Many organs, including the vascular and digestive systems, are immature or absent in the larvae but become functional in the juvenile stage (Karaiskou 2015). Thus, many organs functional in vertebrate larvae are established during ascidian metamorphosis with *Hox* gene involvement, suggesting that certain embryonic events in vertebrates may correspond to post-embryonic events in ascidians.

The collinearity of expression, function, and genomic organization is a central feature of *Hox* cluster genes (Gaunt 2018; Krumlauf 2018). In this context, the suggested role of *Hox2* in the atrial siphon is consistent with this collinearity. Though the ascidian atrial siphon functions posteriorly in the digestive system, serving as an excretory opening, its primordia are morphologically anterior, located in the larval trunk. This structure is also considered homologous to the vertebrate otic placode (Mazet et al. 2005; Wada et al. 1998). Therefore, the dependence of atrial siphon formation on the anterior-class *Hox* gene *Hox2* is consistent with functional studies in vertebrates. *Hox1* also contributes to atrial siphon formation, acting in the epidermis to establish atrial siphon primordia during the larval stage (Sasakura et al. 2012). Both *Hox1* and *Hox2* are epidermal factors, but *Hox1* is required earlier than *Hox2*, suggesting temporal collinearity between them. *Ciona*’s posterior *Hox* genes (*Hox10*, *Hox12*, and *Hox13*) are responsible for forming the intestine, a posterior digestive structure (Kawai et al. 2015; Tajima et al. 2020; Yoshida et al. 2017). Although the tunicate *Hox* cluster is genomically disrupted, its functional collinearity is maintained, indicating that genomic architecture is not essential for this feature (Ikuta et al. 2004; Seo et al. 2004).

The fusion of two atrial siphons is not conserved among tunicates (Satoh 1994). Many ascidian species develop a single atrial siphon from the beginning of the juvenile stage. Atrial siphon formation around metamorphosis follows three patterns: the presence of two atrial siphon primordia in the juvenile stage is limited to *Ciona* and related species. Fusion of two primordia before hatching characterizes the order Aplousobranchia, while a single primordium/siphon is found in the order Stolidobranchia. As *Hox2* is conserved across tunicates (DeBiasse et al. 2020; Sekigami et al. 2017; Seo et al. 2004; Zhang et al. 2021), future studies should examine whether *Hox2* regulates atrial siphon formation at earlier developmental stages in Aplousobranchia and Stolidobranchia than in *Ciona*.

Epidermal cell flattening along the anterior–posterior axis correlates with atrial siphon fusion (Figure 5f-g, Figure 7a). During fusion, the angle between siphons decreases, drawing their edges toward the midline (Ohta et al. 2010). We suspect that epidermal cell flattening drives this angular reduction. The decrease in surface area between two protrusions generates a force pulling them together, akin to pinching the junction between two air-filled fingers of a rubber glove (Figure 7b). This flattening is *Hox2*-dependent. Further research must characterize downstream genes of *Hox2* to elucidate its regulatory mechanisms in promoting atrial siphon fusion. Genes involved in actin dynamics and planar cell polarity are likely to be responsible for this process.

**FIGURE 7.**
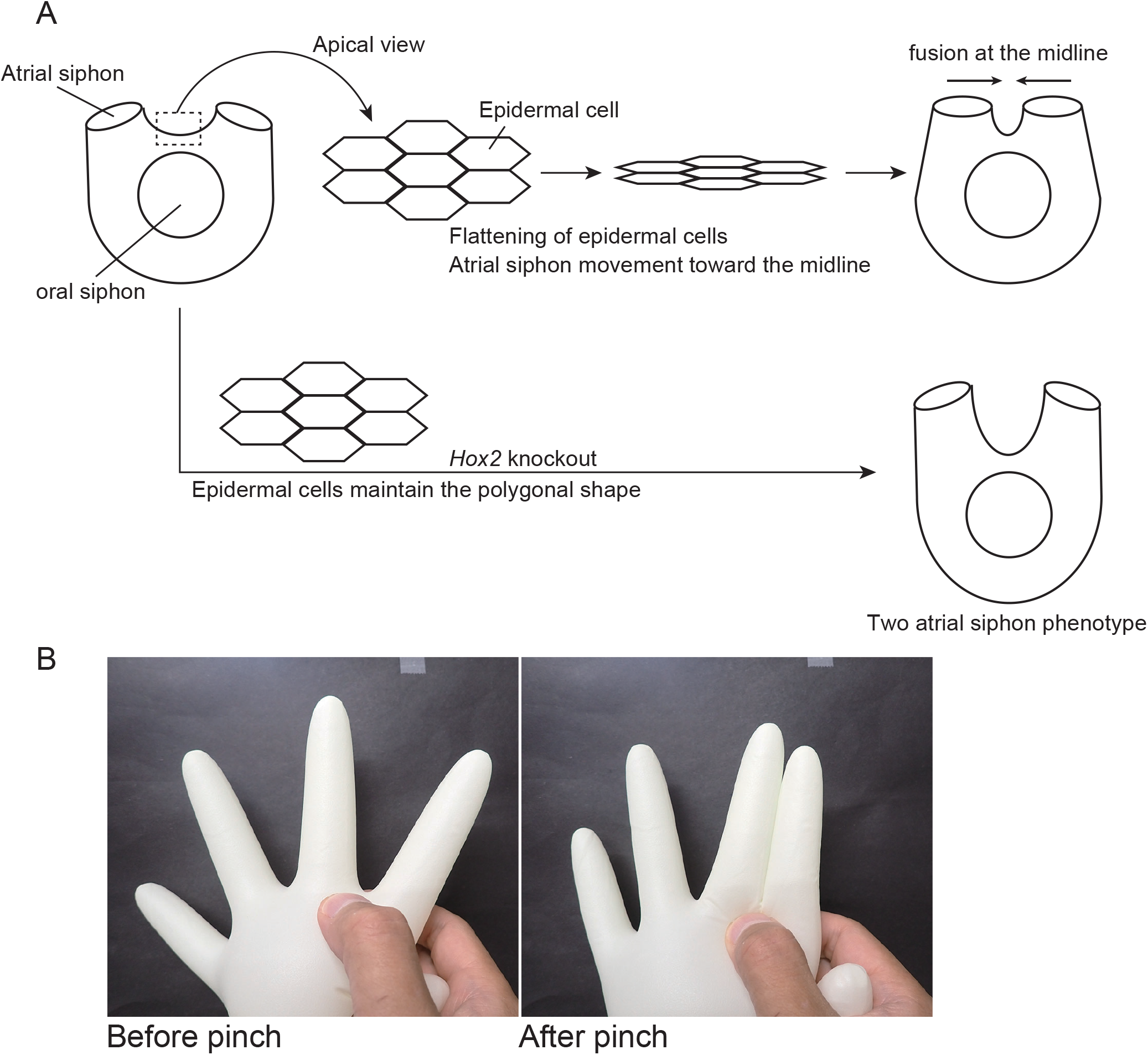
Hypothetical processes of atrial siphon fusion and possible contributions of *Hox2*. (a) Schematic illustration of the relationships among epidermal cell flattening, *Hox2*, and atrial siphon fusion. (b) Two glove fingers joined by pinching their junction.

A limitation of our study was our failure to reproduce two siphon phenotypes using genome-editing constructs other than TALENys. Our data indicate that the *Hox2* locus is the only reproducibly mutated candidate associated with this phenotype, supporting a causal role for TALENys-induced *Hox2* mutations. However, homozygous *Hox2* mutants, which are expected to express a truncated Hox2 protein lacking the homeodomain’s central region, did not exhibit the abnormality. This suggests the two-atrial-siphon phenotype does not arise simply from loss of *Hox2* function. One possibility is that other Hox proteins compensate for *Hox2* function in the mutants. This possibility is supported by the nature of Hox proteins, which recognize almost identical AT-rich nucleotide stretches (De Kumar & Darland 2021).

TALENys-introduced G0 animals exhibited mosaic mutations, raising the possibility that a specific mutation causes the phenotype. These TALENys-induced mutations include base deletions in multiples of three, resulting in the deletion of a few amino acids without altering the protein’s major reading frame (Figure 2a). Such subtle amino acid changes may yield proteins with altered functions, such as modified cofactor affinities. These abnormal proteins might act as dominant-negative forms, strongly suppressing transcription of downstream genes essential for the fusion of atrial siphons. A phenomenon that may be related to *Ciona Hox2* has been observed with mammalian *Hoxd13* (Bruneau et al. 2001; Goodman et al. 1997; Wang et al. 2024). For example, the Hoxd13 Q50R mutation is pathogenic in syndactyly type V, causing severe malformation of the fourth and fifth metacarpals. Heterozygous Hoxd13(Q50R) exhibits abnormality, indicating the mutation overwhelms wild-type Hoxd13 protein function. Similarly, a specific point mutation in the homeodomain of *Ciona* Hox2 may dominantly suppress the fusion of atrial siphons. Future studies characterizing the causative *Hox2* allele and its downstream genes will clarify *Hox2*’s role in the two-atrial-siphon phenotype.

## ACKNOWLEDGMENTS

We thank Dr. Yutaka Satou for helpful discussions regarding genome analyses and for providing wild-type *Ciona*. We are grateful to the Shimoda Marine Research Center at the University of Tsukuba for animal maintenance, and to Drs. Shigeki Fujiwara, Manabu Yoshida, and all members of the Department of Zoology, Kyoto University, the Misaki Marine Biological Station, the University of Tokyo, the Maizuru Fishery Research Station of Kyoto University, and the National BioResource Project (NBRP) for cultivating and providing *Ciona* adults and experimental materials. This study was supported by Japan Society for the Promotion of Science grants to YS (16H04815, 19H03262, 25K02306). YS also received support from a Takeda Bioscience Research Grant.

## Notes

### Competing Interest Statement

The authors have declared no competing interest.

